# Reduced representation optical methylation mapping (R^2^OM^2^)

**DOI:** 10.1101/108084

**Authors:** Assaf Grunwald, Hila Sharim, Tslil Gabrieli, Yael Michaeli, Dmitry Torchinsky, Matyas Juhasz, Kathryn R Wagner, Jonathan Pevsner, Jeff Reifenberger, Alex R Hastie, Han Cao, Elmar Weinhold, Yuval Ebenstein

## Abstract

Reduced representation methylation profiling is a method of analysis in which a subset of CpGs is used to report the overall methylation status of the probed genomic regions. This approach has been widely adopted for genome-scale bisulfite sequencing since it requires fewer sequencing reads and uses significantly less starting material than whole-genome analysis. Consequently, this method is suitable for profiling medical samples and single cells at high throughput and reduced costs. Here, we use this concept in order to create a pattern of fluorescent optical methylation profiles along individual DNA molecules. Reduced representation optical methylation mapping (R^2^OM^2^) in combination with Bionano Genomics next generation genome mapping (NGM) technology provides a hybrid genetic/epigenetic genome map of individual chromosome segments spanning hundreds of kilobase pairs (kbp). These long reads, along with the single-molecule resolution, allow for epigenetic variation calling and methylation analysis of large structural aberrations such as pathogenic macrosatellite arrays not accessible to single-cell next generation sequencing (NGS). We apply this method to facioscapulohumeral dystrophy (FSHD) showing both structural variation and hypomethylation status of a disease-associated, highly repetitive locus on chromosome 4q.

## INTRODUCTION

DNA methylation, specifically methylation of the 5-carbon of cytosine, is the most studied and among the most significant epigenetic modifications^1^. In mammalian DNA, methylation mostly occurs on cytosines within CpG dinucleotides (DNA motifs where the cytosine is followed by a guanine residue) that are methylated to an extent of ~70%^2^. Approximately 60% of human gene promoters contain clusters of CpGs referred to as CpG islands (CGIs)^3^. CpG methylation plays an important role in regulation of gene expression with the general notion that hypermethyation of promoters suppresses gene expression^4^. Thus, the methylation status of gene promoters may predict gene activity and relate epigenetic transformations in development and disease to gene expression and protein abundance.

Another genetic feature regulated by CpG methylation is repetitive arrays. These variable copy number elements are homologous DNA sequences, exhibiting identity (or great similarity) to each other^5,6^. Many repetitive elements are mobile and can transpose across the genome, perform homologous recombination events, and promote dynamic genomic transformation^6–8^. This unstable nature explains their size variability, both among different individuals and between different cells of the same individual^6,9,10^. Typically arrays are characterized by the number of units composing them, which has been shown to affect their activity^11^. Methylation of repeat units adds another dimension of variability to these elements: Locally it may regulate the function of individual units and globally it can change the effective number of units in an array altering its activity. In this context, it has been shown that methylation levels of repetitive DNA can regulate repeat-related genetic diseases^12,13^, and are correlated with various types of cancer and their severity^14^. One striking example of a repeat array, addressed in this work, is facioscapulohumeral muscular dystrophy (FSHD), one of the most commons form of muscular dystrophy, affecting approximately 1 in 7,000-20,000 individuals^15–18^.

The "gold-standard" method for studying CpG methylation is bisulfite sequencing, often used for whole-genome, base-pair resolution methylation profiling by next generation sequencing (NGS). As such, it provides an averaged representation of the sample’s DNA sequence, with distinction between methylated and non-methylated cytosines^19^. The main shortcomings of bisulfite sequencing are the high cost and the requirement for large amounts of starting material due to chemical degradation of the DNA, and the need to split the sample for sequencing the treated and untreated samples separately.

Reduced representation bisulfite sequencing (RRBS) utilizes restriction enzymes that cut at CpG sites, thus specifically enriching for fragments that end within CpG islands. The fragment ends are ligated to adapters, size selected, bisulfite converted, and sequenced to give a representative genome-scale methylation profile. Most commonly, the restriction enzyme MspI is used to cut DNA at CCGG sites. The NGS output captures 60% of gene promoters, thus producing important regulatory information while requiring very little input sample^20^. The low input implies that fewer reads are required for accurate sequencing, allowing for high throughput, low-cost methylation analysis for clinical and single-cell applications^21,22^. Nevertheless, bisulfite sequencing and RRBS have limited accessibility to repetitive regions due to the inherent limitations of short-read sequencing, and are often unable to quantify the length and arrangements of the repeats ^23^ (supporting information). Thus, in many cases the genetic and epigenetic characteristics of repetitive elements are still unknown. With this in mind, it is plausible that some of the many NGS-inferred sequence duplications are in fact longer repetitive elements. Moreover, since both repetitive elements and methylation patterns tend to display somatic mosaicism, manifesting different structures in different cells^10^, their variability is often masked in the averaged NGS results.

Here we harness the reduced representation concept in order to access the methylation profile of long individual chromosome segments by optical genome mapping. Reduced representation optical methylation mapping (R^2^OM^2^) seeks to simultaneously capture large scale structural and copy number variants together with their associated methylation status.

DNA optical mapping^24–28^ stands out as an attractive approach for studying large genomic rearrangements such as repeat arrays. The approach consists of a set of techniques for stretching long genomic fragments, followed by imaging of these fragments using fluorescence microscopy. Image processing is used to read out a labeled specific sequence-motif pattern physical barcode along the molecules that provides genetic information such as the genomic locus of the imaged molecule as well as the size and number of large repeat units. The most advanced method for optical genome mapping involves linearizing and uniformly stretching of fluorescently barcoded DNA molecules in highly-parallel nanochannel arrays. This technique, commercialized by Bionano Genomics Inc., is capable of large scale genetic mapping and automated copy number analysis on a genome scale^29,30^.

The use of fluorescence microscopy presents the potential of obtaining, in parallel and from the same molecule, several types of information by labeling different genomic features with different colors. This allows access to epigenetic marks and DNA damage lesions at their native genomic context on the single molecule level^31–33^. In order to facilitate such multiplexing, we have developed an enzymatic labeling reaction that can distinguish methylated from nonmethylated cytosines, and showed that it can be used to detect and quantify methylation levels in synthetic DNA molecules translocated through solid-state nanopores^34^. The bacterial methyltransferase M.TaqI in combination with a synthetic cofactor analogue can fluorescently label the adenine residue in TCGA sites containing non-methylated CpGs. M.TaqI has potential access to only about 5.5% of all CpGs, however, these sites represent over 90% of gene promotors and hence M.TaqI is a good candidate for reduced representation methylation profiling for genomic regulatory function study. Combining this profile with a genetic pattern generated by labeling of specific sequence motifs, results in a hybrid genetic / epigenetic barcode in the same field of view. Using the genetic labels for mapping to the sequence reference allows, for the first time, simultaneous assignmentof the epigenetic labels to their specific genomic locations, and the study of DNA methylation patterns over large genomic fragments at single genome resolution.

We compare the methylation levels of primary white blood cells and a B-Lymphocyte derived cell line [NA12878], directly quantifying the reduction in CpG methylation levels after Epstein-Barr virus cell-line transformation. Furthermore, we show that optical methylation profiles are correlated with documented sequencing results and that optical signal levels from promotor methylation islands correlate with published gene expression data^35,36^. Finally, we demonstrate the utility of R^2^OM^2^ to simultaneously probe copy number and methylation level in microsatellite arrays, that when contracted can result in disease, serving as a potential diagnostic assay for FSHD.

## RESULTS

### DNA methylation detection at the single-molecule level by DNA-methyltransferase assisted labeling

DNA methyltransferase enzymes (MTases) offer an orthogonal method for DNA sequence-specific labeling as they may be "tricked", using synthetic cofactor analogs, into directly incorporating a fluorophore onto their recognition site^37,38^ (Figure 1a upper panel). We recently demonstrated how the DNA MTase M.TaqI (with the TCGA recognition sequence) can generate DNA barcodes for optical mapping and bacteriophage strain typing by transferring a fluorophore onto the target adenine in its recognition site^39^. However, the M.TaqI recognition site contains a nested CpG dinucleotide, which when cytosine-methylated blocks adenine methylation by M.TaqI^40^. To test whether this property of M.TaqI could also be used to label DNA depending on the CpG methylation status, we first used a bulk restriction-protection assay (Figure 1b and supporting information). The assay utilizes the R.TaqI restriction enzyme that specifically cleaves DNA at TCGA sites containing unmodified adenine. Consequently, digested DNA confirms the inability of M.TaqI to label the adenine when the CpG within the recognition site is methylated. Additionally, the assay shows a 100% labeling efficiency even at a 1:64 ratio of M.TaqI to labeling sites, emphasizing the catalytic performance of this DNA MTase enzyme. We further tested the specificity of the labeling reaction at the single molecule level (Figure 1c). 50 kbp λ-bacteriophage genomes, containing 121 M.TaqI recognition sites, were labeled with the fluorescent cofactor AdoYnTAMRA, yielding a unique continuous fluorescent signal along individual genomes that were deposited and imaged on a microscope slide. After methylating all CpGs on the λ-fbacteriophage genomes by *in-vitro* treatment with the CpG MTase M.SssSI, essentially no labelling was detected on the deposited λ DNA, corroborating the bulk results that the labelling reaction is completely blocked by existing CpG methylation.

**Figure 1.**
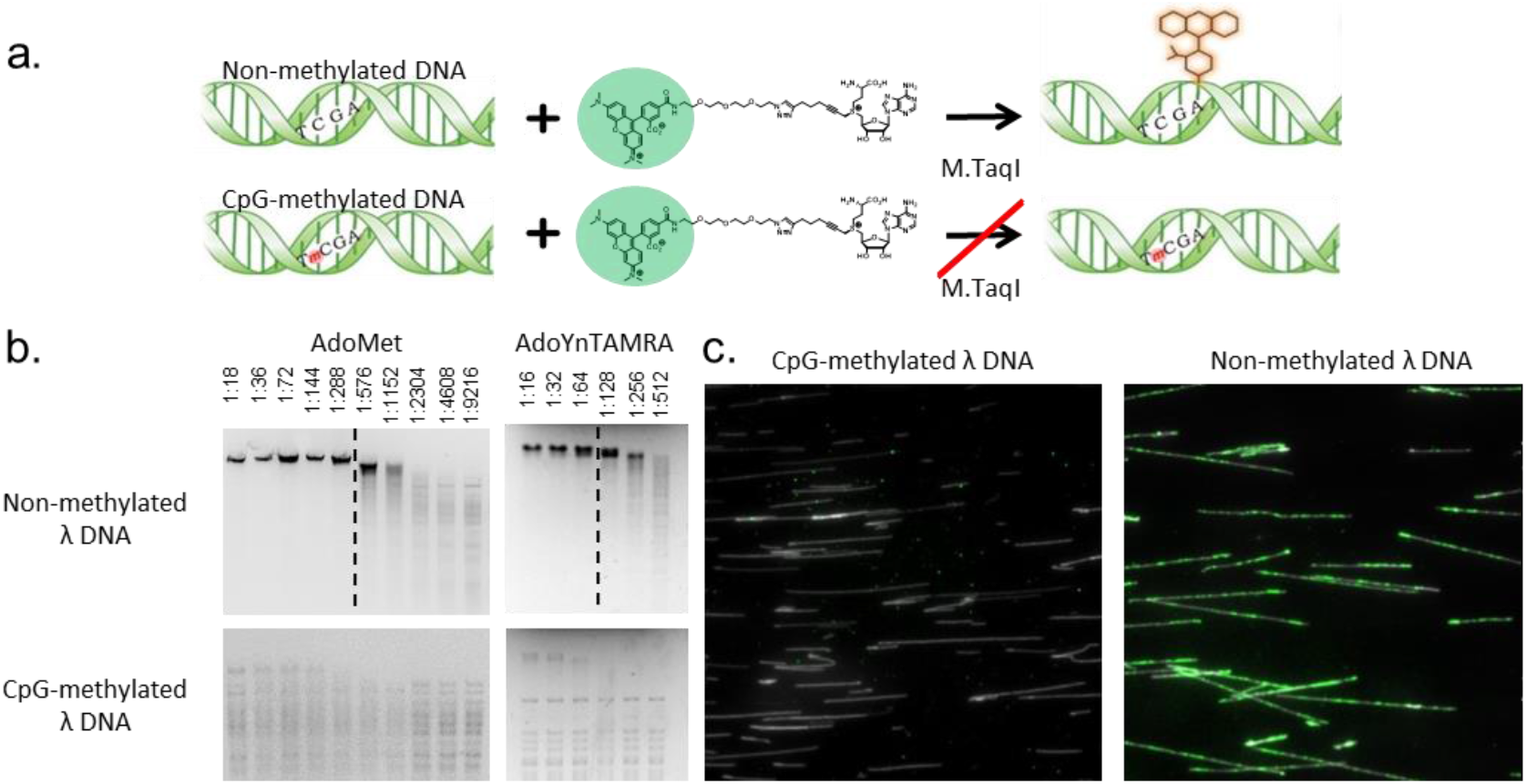
**a**. Top: M.TaqI catalyzes the transfer of a TAMRA fluorophore from the cofactor AdoYnTAMRA onto the adenine residue that lies in its TCGA recognition site. Bottom: If the cytosine residue that lies within M.TaqI’s recognition site is methylated, the reaction is blocked. **b.** Gel images from "protection assays" that were performed to confirm M.TaqI sensitivity to cytosine methylation. Either non-methylated or CpG-methylated λ DNA was reacted in the presence of the natural cofactor AdoMet (left panel) or the synthetic fluorescent cofactor AdoYnTAMRA, (right panel) using decreasing amounts of M.TaqI (enzyme per sites ratio indicted above the lanes). Following, the DNA was challenged with the adenine methylation-sensitive restriction enzyme R.TaqI. Finally, the reactions were size separated using gel electrophoresis. From the resulting gel images, it is clear, for both cofactors, that while non-methylated DNA is modified by M.TaqI and thus protected from R.TaqI restriction (up to a critical enzyme to sites ratio from where fragmentation is observed, represented by a dashed line). CpG-methylated DNA, does not get modified and thus restriction is observed at all M.TaqI concentrations, for both cofactors. **c.** Non-methylated or CpG-methylated λ DNA was reacted with M.TaqI and the fluorescent cofactor AdoYnTAMRA, the DNA’s backbone was labeled with YOYO-1. Labeled DNA was stretched on modified glass surfaces and imaged in two channels to visualize its contour (displayed in grey in the images) and M.TaqI fluorescent labelling (displayed in green). While the methylated DNA remained unlabeled the non-methylated DNA was heavily labeled.

### Optical mapping quantifies genome-wide and locus specific methylation levels

We next tested whether the methylation sensitive labeling reaction may be combined with optical mapping, for studying the methylation levels of various genomes and specific DNA regions. We chose to compare primary blood cells to an Epstein-Barr virus (EBV) transformed blood lymphocyte cell-line, NA12878 (Coriell Institute for Medical Research, Camden New Jersey, USA), where reduced genome-wide methylation levels have been reported^41^.

Genomic DNA was first nick-labeled with the nicking enzyme Nt.BspQI, to create a distinct genetic pattern of red labels along the DNA, regardless of the epigenetic state of the sample^42^. Next, a second layer of information was added by labeling non-methylated CpGs with a green fluorophore using M.TaqI. The labeled DNA was loaded into nanochannel array chips, for dualcolor optical genome mapping (Figure 2a). We used the IrysView software suite (Bionano Genomics Inc.) for automatic label detection and genomic alignment of the imaged molecules. Our goal was to provide global quantification of the relative methylation levels in the two samples. As DNA extracted from EBV immortalized cell lines is expected to be hypomethylated compared to DNA from primary cells^41^, a higher number of M.TaqI labels was expected for this sample. Indeed, while the number of genetic labels was similar for both samples, the cell line DNA displayed more labelling in the epigenetic channel (16.96 labels per 100 kbp) compared to the primary DNA (4.73 labels per 100 kbp) as predicted (Figure 2b). Since close-by CpGs are unresolved due to the limits of optical resolution, we also analyzed the intensity of the methylation labels in order to gain information regarding the number of labeled CpGs. Such analysis shows an increase in the methylation level ratio between the samples from 3.6 to 6.3 (Figure 2b). These results demonstrate the utility of M.TaqI labeling for quantitative assessment of methylation levels in various samples.

**Figure 2.**
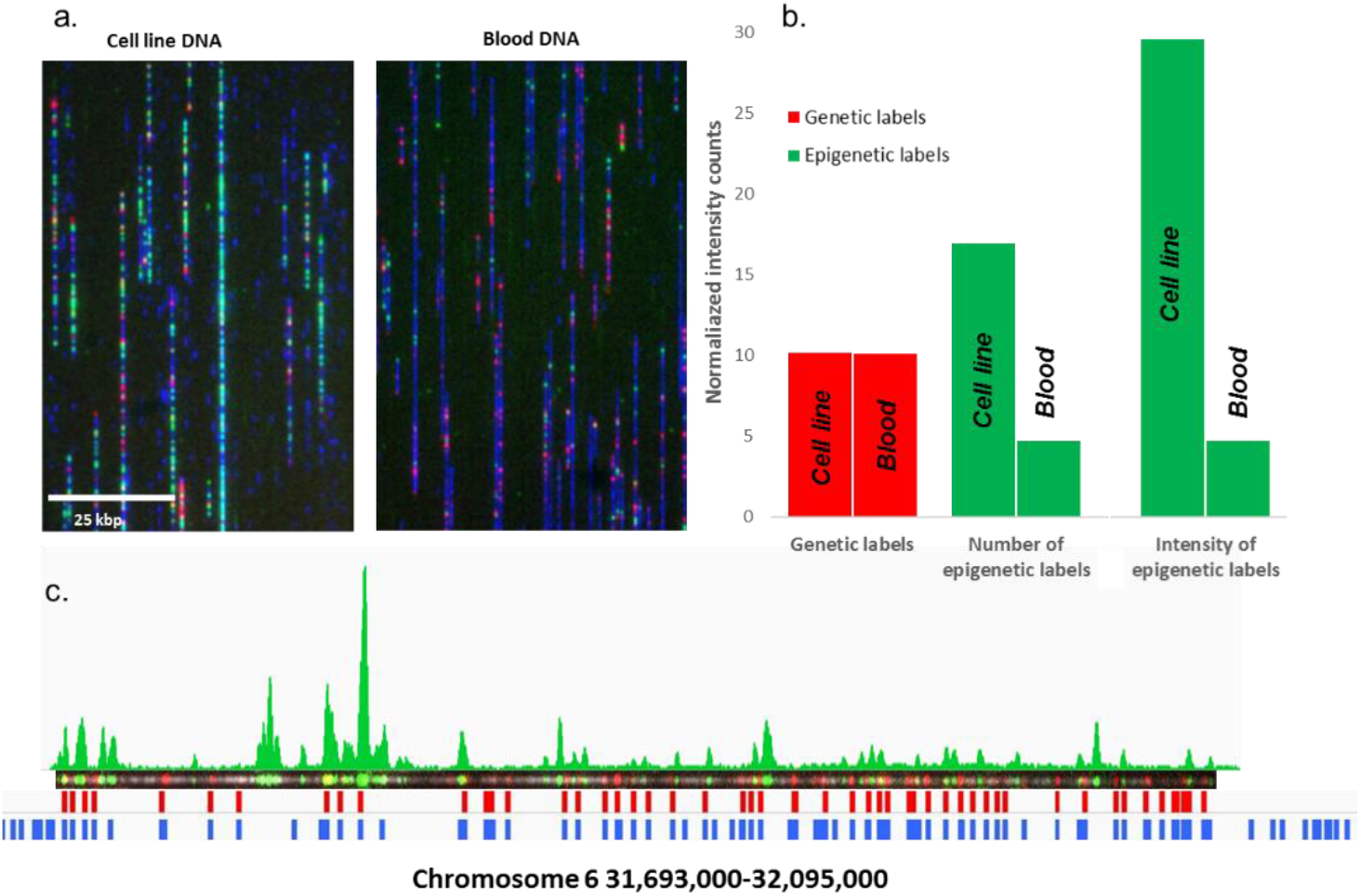
Methylation detection on genomic DNA. **a.** Genomic DNA from both primary human blood cells (right image) or a human lymphocyte cell line (left image) was dual labeled with genetic labels (Nt.BspQI, red labels) and methylation sensitive labels (M.TaqI, green labels), and stretched and imaged on a nanochannel array chip. Representative images from both samples are presented here. DNA backbone, stained with YOYO-1 is displayed in blue. **b.** Genetic and epigenetic labels were detected for both DNA samples. The bar-graph presented here displays number of detected labels per 100 kbp. The genetic labels are displayed in red bars and the epigenetic label count in green. The two right most bars display normalized intensities of epigenetic labels per 100 kbp in both samples. It is clear that while genetic labels were detected at similar amounts in both samples, epigenetic labels are more frequent in the immortalized cell line genome, indicating that it is highly non-methylated as expected. **c.** A representative dual labeled molecule from the primary blood sample. Epigenetic label intensity distributions of labels along the molecule are presented above the molecule image. The detected locations of genetic labels are shown in red below the image, and the reference barcode, used for alignment of the genetic labels is shown in blue.

To study local methylation patterns at specific genomic locations, the genetic barcode was used to align the molecules to their corresponding sequence locations. After alignment, the epigenetic maps were used to study methylation patterns along individual DNA molecules as shown in Figure 2c. for a 400 kbp molecule mapped to chromosome 6.

In order to demonstrate the ability of R^2^OM^2^ to detect epigenetic variation and predict gene expression via DNA methylation mapping, we examined three DNA molecules from primary white blood cells with ~250 kbp genetic overlap. Figure 3 displays published genomic methylation and gene content data as well as the individual molecular methylation maps at a specific genomic region on chromosome 1. The upper panels show the distribution of CpG and M.TaqI sites in this region as well as recently published white blood DNA methylation data obtained by MeDIP sequencing^43^. We note that labelling is expected at non-methylated CpGs and use the sequencing data to predict labeling at M.TaqI sites overlapping non-methylated regions (green line marks in Figure 3c). The molecules in Figure 3d exhibit similar methylation profiles that are well correlated with the expected population level sequencing data. Small variations in the pattern are seen and can be attributed to common variation in methylation status among white blood cells^44^. Furthermore, the maps allow relating promoter methylation to gene expression data. For instance, a CpG island that lies in the promotor region of the *PHF13* gene (indicated by a red box), was found to be non-methylated by both sequencing and R^2^OM^2^, and is known to be expressed in white blood cells^36^. In contrast, the promoter of the *KLHL41* gene (indicated by a black box), was detected as methylated by both methods. This muscle specific gene is indeed expected to be silenced in blood cells^35^. These findings and their agreement with published results obtained by NGS, validate the ability of R^2^OM^2^ to unravel the methylome at a single genome level. Optical mapping allows genomic information to be read directly from individual, un-amplified, long fragments of DNA, thus enabling comparison of methylation patterns between genomes across hundreds of kbp, mitigating the bias of large structural and copy number variation in single cell sequencing experiments.

**Figure 3.**
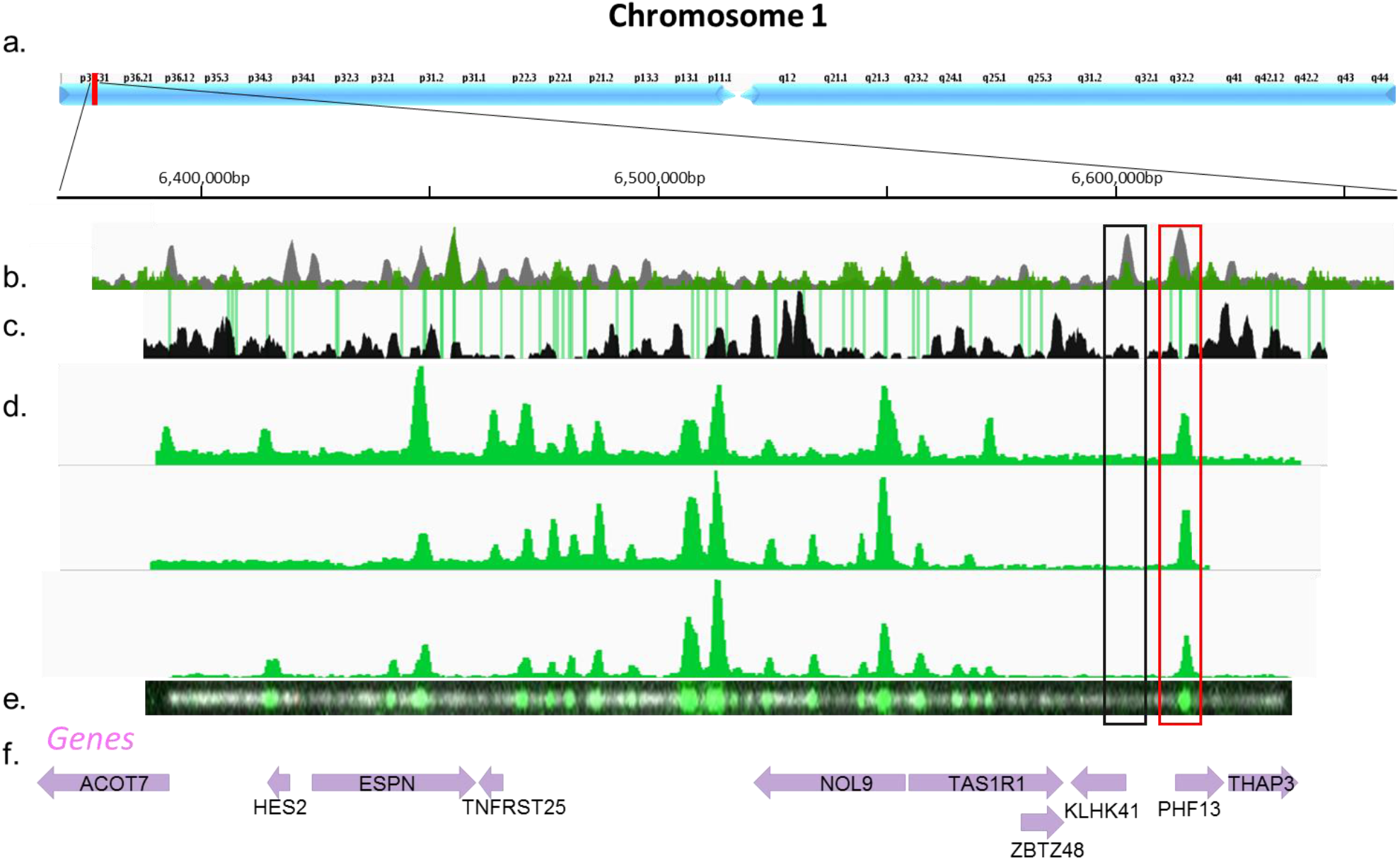
**a.** Detailed view of chromosome 1:6,350,000-6,670,000 bp. **b.** A histogram of CpG (gray) and M.TaqI (green) sites across the region. **c.** A coverage histogram of white blood cells MeDIP-seq. Peaks indicate high methylation levels, and valleys indicate low methylation levels - where labeling is expected. The green lines represent M.TaqI recognition sites overlapping with areas where non-methylated CpGs are expected from the MeDIP-seq. These are the predicted sites to be labeled by M.TaqI. **d.** Digital representation of three detected molecules aligned to the reference sequence based on Nt.BspQI labels (P-value<10^−20^). Label intensity is shown in green. **e.** The raw image of the backbone (light gray) and the M.TaqI channel (green)of the bottom molecule presented in d. **f.** Gene body locations and corresponding HGNC gene symbols. Each gene is represented as a purple arrow and the direction of the arrow indicates gene orientation. Black and red rectangles across b-e indicate methylated and non-methylated gene promoters, respectively

### Simultaneous quantification of copy number and methylation state in DNA tandem repeats

Macrosatellite arrays, repetitive DNA that spans up to millions of base pairs across the genome, are extremely challenging for analysis by next-generation sequencing. Analysis is further complicated by the recent understanding that DNA methylation plays a crucial role in the function of these regions. One striking example of the significance of methylation of such regions is the D4Z4 array on chromosome 4, which is directly related to Facioscapulohumeral muscular dystrophy (FSHD)^45^. It has been shown that both the number of D4Z4 repeats and their methylation status constitute the genotype of the disease, dictating whether the individual has weakness or not. Commonly, healthy individuals carry an array of more than 10 repeats. However, even long arrays result in FSHD symptoms when losing their methylation (termed FSHD2) while carriers of short but highly methylated repeat arrays, do not manifest the disease^15,16^. The multiple combinations of copy number and methylation level span a broad range of possible variations which are correlated with the disease severity and manifestation ^16,17^.

To demonstrate the capabilities of optical mapping for simultaneous copy number quantification and DNA methylation detection, we studied a model system of the FSHD-associated D4Z4 repeats cloned into the CH16-291A23 bacterial artificial chromosome (BAC) (Figure 4a). We first attempted to evaluate the copy number of the studied array by NGS read-depth analysis^46^. We reasoned that the number of reads representing the repeat unit, relative to the number of reads detected for a single copy region, would provide the array copy number. Purified BAC DNA was sequenced using Illumina MiSeq to a read depth of 15,000X. We used read depth analysis to assess the D4Z4 copy number. Briefly, all sequencing reads were aligned to the BAC’s reference sequence, containing only one repeat, and the copy number was calculated as the ratio between the number of reads ("coverage") aligned to the repetitive and the non-repetitive regions respectively. The coverage along the non-repetitive sequence displayed variation of up to 25%, while the coverage along the repeat sequence was extremely variable, averaging 63% of the mean read depth (Figure S1). The ratio between the median coverage value along the repetitive and non-repetitive sequences was ~8, implying that this is the number of D4Z4 repeats along the BAC (Figure 4c). However, the large standard deviation values strongly demonstrate the unreliability of this method, as well as the sensitivity of PCR amplification and NGS to the exact content of the investigated sequence.

**Figure 4.**
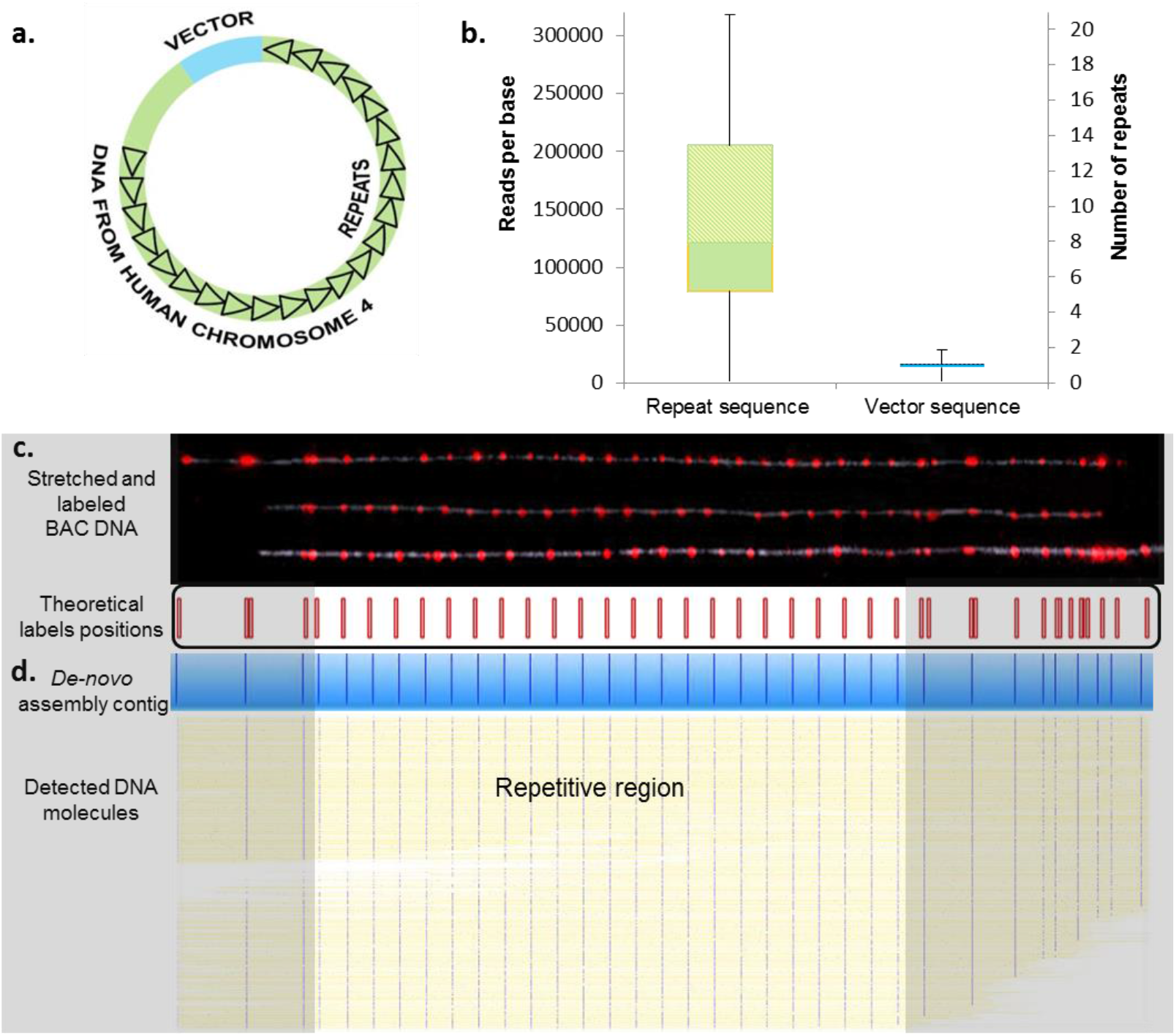
Copy number analysis. **a.** The FSHD BAC model system contained an unknown number of D4Z4 repeats (black triangles), an unknown size of genomic DNA 5’ to the repeat array (green), and the cloning vector (blue). **b.** Read depth analysis box plots displaying the 25th percentile (bottom of the lower box), the median (intermediate between boxes) and the 75th percentile (top of the upper box), for coverage values of the sequencing reads along the repeat region (left box), and the non-repetitive region (including the vector and the upstream to repeat region, right box). The scale on the right is normalized to the median coverage along the non-repetitive region. **c.** Representative images of three intact model molecules labeled with Nb.BsmI (red dots) and stretched on modified glass surfaces. The pattern of labeled molecules can be aligned to the reference map presented below the images (expected labeling locations are shown in red). The repetitive region can be distinguished by virtue of the equally spaced labels, each representing a repeat, and quantified by simply counting the labels (23 repeats). **d.** 627 digital representations of labeled DNA molecules stretched and imaged in a nanochannel array chip. The detected molecules were *de-novo* assembled into a single consensus map (displayed in blue) which shows excellent agreement with the theoretical map. The yellow lines represent the DNA backbone and the blue dots represent the detected labels. The assembled map contained 23 repeat units.

Next, we directly visualized the repeats along stretched DNA molecules. We reasoned that tailored labeling of specific sequence motifs would highlight individual repeats and allow for physical counting of the copy number. We nick-labeled the BAC DNA using nicking enzymes (Nb.BsmI or Nb.BssSI) which have a single recognition site on the 3.3 kbp long D4Z4 repeat sequence, yielding a single distinct fluorescent tag for each repeat unit. The labeled DNA was stretched and immobilized for visualization on modified glass surfaces, using a simple microfluidic scheme. This method allowed fluorescent imaging of the entire DNA contour and localization of individual fluorescent labels along the DNA (Figure 4c). The same sample was loaded onto an Irys instrument (Bionano Genomics Inc., CA. USA), which facilitates high-throughput DNA stretching and imaging in nanochannel array chips. The post-imaging analysis, performed by the IrysView software suite, involved automatic label detection and *de-novo* assembly of the molecules into a continuous consensus barcode. The resulting consensus map was created in an unsupervised manner based on label patterns from approximately one thousand detected molecules^47,48^. When comparing the non-repetitive region of this contig to the one predicted from the known sequence, an almost perfect match was obtained (p-value <10^−43^, figure 4d). The consensus repetitive region is unambiguously composed of 23 D4Z4 repeats.

For R^2^OM^2^ we performed M.TaqI labeling in order to overlay the epigenetic map on the repetitive genetic barcode. We used red fluorophores for the genetic barcode and green fluorophores for methylation mapping. On the non-methylated BAC, M.TaqI creates a unique pattern along the non-repetitive region and has two close-by recognition sites on each repeat unit (resulting in one visual label on each repeat, due to diffraction limits)^49^ (Figure 5a). As expected, the non-methylated BAC DNA was dually labeled by both enzymes and thus contained two barcode layers, in agreement with the theoretical dual-color barcode (Figure 5b).

**Figure 5.**
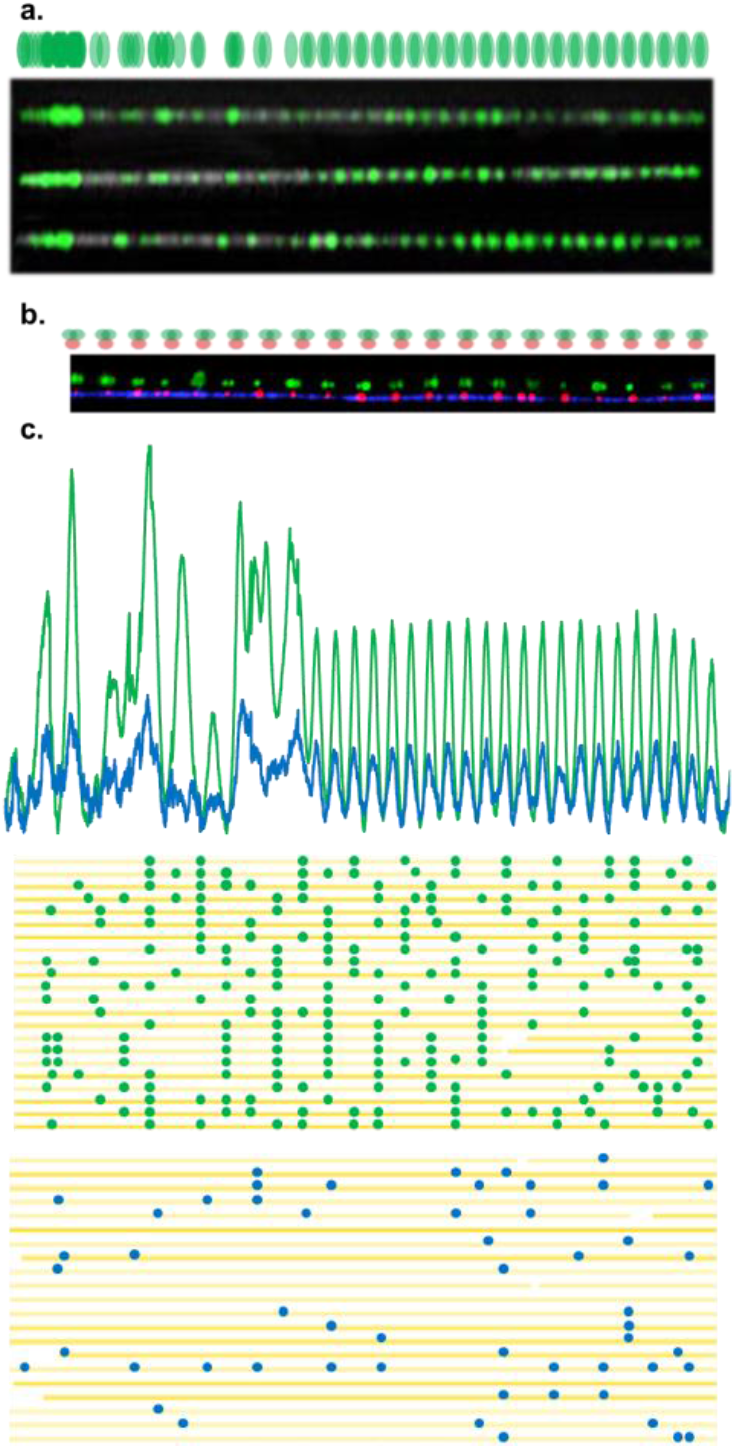
**a.** A reference map simulating the relative expected locations of R^2^OM^2^ labels generated by the MTase M.TaqI along the stretched BAC (green). Below, images of three non-methylated, M.TaqI labeled BAC molecules linearized and stretched in nanochannels and aligned to the reference map. **b.** R^2^OM^2^ of a region from FSHD BAC stretched on a modified glass slide. The genetic identity and the number of repeat units are highlighted by labeling with the nicking enzyme Nb.BsmI (red labels). Co-labeling the DNA with M.TaqI (green labels) indicated that the molecules were non-methylated. The displayed image is an overlay of red and green channels along the repetitive region of a single BAC molecule. Above the repeats is the reference map for the region. The green channel is shifted upwards to allow better visualization. The higher stretching factor achieved on modified glass compared to channels, enables detection of the two M.TaqI labels flanking the genetic label in each repeat. **c.** Comparative R^2^OM^2^ of non-methylated and partially-methylated BAC samples. Normalized integrated maps of detected M.TaqI labels are presented for the non-methylated (green, 18074 molecules) and partially-methylated (blue, 9089 molecules). Both plots highly correlate with the expected reference map. Digital representation of several nonmethylated (top, green labels) and methylated (bottom, blue labels) molecules. The backbone of the DNA is shown in yellow and detected M.TaqI labels in green/blue.

### High-throughput, automatic detection of variable methylation levels at single-repeat resolution

To simulate the native state of DNA, where repeat arrays are methylated to variable degrees, we partially methylated the DNA using the CpG-specific DNA MTase M.ssSI. We repeated the dual-labeling reaction on the partially methylated sample, as well as on a non-methylated control, and analyzed both on the nanochannel array chips. After image analysis, we used the red genetic labels for automated *de-novo* assembly and generated the consensus map with the 23 repeats as described earlier (and depicted in Figure 4d). With thousands of molecules now aligned to the consensus map, we could compare the green methylation patterns generated by R^2^OM^2^ on the non-methylated and partially methylated samples. Figure 5a shows three representative fluorescent methylation maps that are clearly in agreement with the theoretical reference for M.TaqI along the BAC. The non-methylated sample shows frequent labeling, at the expected locations, while the partially methylated DNA sample shows significantly fewer labels (Figure 5b). To further compare the two samples, we generated plots of integrated label counts based on thousands of molecules detected in each data set (Figure 5c). The amount of labeling on the non-methylated model was significantly higher than that along the methylated BAC (p-value < 10^−50^, t-test). It is clear from the plot that in the partly methylated sample, methylated CpGs are distributed equally among the repeat units, in line with the fact that the partial methylation was random and equal for all repeats in the array. These results clearly demonstrate that R^2^OM^2^ enables not only single-molecule and single-repeat resolution but also assessment of the average methylation status for each individual repeat in the array across a population of different DNA molecules.

Finally, we performed R^2^OM^2^ on a FSHD patient sample (Figure 6). DNA was extracted from whole blood of a donor previously diagnosed via commercial testing based on pulse field gel electrophoresis and Southern blot. DNA was labeled and analyzed on the Irys instrument. Genetic labeling (marked red) allowed distinguishing between the two very similar 4qA and 4qB alleles and the microsatellite repeats are visually marked by the equally spaced methylation sensitive labels (marked green) indicating a non-methylated array. We counted 8 repeat blocks on the non-pathogenic 4qB allele and 5 repeat blocks for the 4qA allele^50^. This corroborated previous genetic assessments based on clinically performed pulsed-field gel electrophoresis in which the 4qB allele was estimated to have a size of 27 kb (EcoRI/BlnI Southern blotting) or 39 kb (EcoRI Southern blotting), corresponding to 8 D4Z4 repeats. These data demonstrate the utility of this optical mapping approach for characterizing the genetic and epigenetic profile of FSHD as well as other microsatellite arrays and large structural variants.

**Figure 6.**
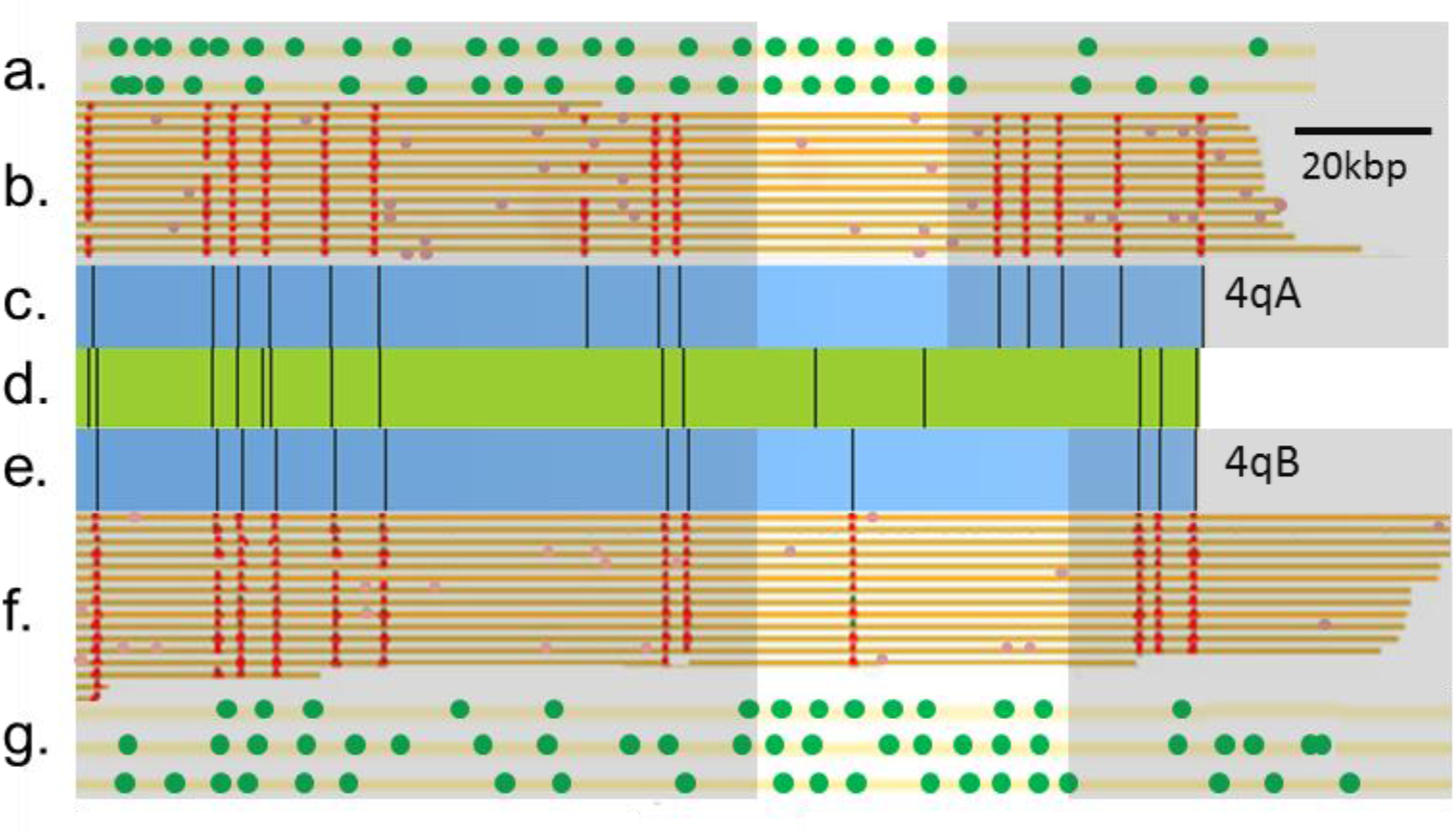
Structural variation analysis and methylation analysis of the pathogenic contraction on chromosome 4q of a patient with FSHD. The chromosome 4 reference map is shown (green horizontal bar, **d.**). The consensus optical maps of the two haplotypes, 4qA (**c.**) and 4qB (**e.**), generated by *De-Novo* assembly are shown. **b./f.** show genetic maps generated from individual molecules aligned to the genome reference. Maps were clustered by haplotype (4qA and 4qB). Red dots represent labels used for alignment. **a./g.** show R^2^OM^2^profiles of DNA molecules aligned to the 4qA/4qB allele and displaying 5 units (4qA) and 8 units (4qB) of the D4Z4 repeat. The gray background covers the non repetitive region and repeats are counted according to the equally spaced R^2^OM^2^ labels which also indicate that the array is hypomethylated.

## DISCUSSION AND OUTLOOK

Optical genome mapping has recently been used to map whole human genomes at high coverage and to highlight genetic variability between individuals with unprecedented detail^29,30^. This work adds an epigenetic component to optical maps, providing, for the first time, correlated genetic/epigenetic profiles for individual DNA molecules spanning hundreds of thousands of base pairs. Utilizing state-of-the-art optical genome mapping technology, combined with DNA MTase-assisted methylation detection, we create a reduced representation optical methylation map, R^2^OM^2^, which reports on the methylation status of 90% of gene promoters. The reduced representation provides sufficiently sparse labeling to comply with the low resolution of optical maps. We demonstrate automatic, high-throughput analysis of global methylation levels in various cell types by quantifying the amount of nonmethylated CpGs. Additionally, R^2^OM^2^ provides long read / low-resolution methylation profiles at different genomic loci with emphasis on CpG islands and regulatory sequence elements such as gene promoters. One area in which R^2^OM^2^ has a particular advantage over existing technologies is the study of DNA methylation of large structural variants and repetitive arrays. DNA repeats are dynamic regions, exhibiting high variability both in length and methylation status, and thus may differ significantly between individuals^9,51^. Today, in the era of personalized medicine, it is becoming increasingly accepted that the full profile of genomic structural variation, including DNA repeats, is directly linked to health and susceptibility to disease^8,52^. R^2^OM^2^ offers single-molecule level information on the size of the region, the number of repeat units, and the methylation status of individual repeats. This detailed information is inaccessible via current genome technologies such as NGS, DNA arrays, and quantitative PCR, which mostly provide averaged or inferred data and cannot specifically address individual repeat units.

We demonstrate the utility of R^2^OM^2^ for characterizing the D4Z4 repeat array, where both the size and the methylation status of the repeats affect FSHD disease manifestation^16^. Specifically, the reported approach can distinguish between healthy individuals, FSHD1, and FSHD2 carriers by combining copy number and methylation level information. Moreover, the detailed view of the methylation status of individual repeats may offer new insights into the mechanism of disease and may lead to a more individualized prognosis than is offered by current commercial testing. Notably, the R^2^OM^2^ concept is not limited to D4Z4 repeats, and various labeling enzymes may be used to address different target sequences, thereby extending the repertoire of available targets for this method. Further development of this technique may serve to map differentially methylated single-molecule patterns on a genomewide scale, potentially allowing simultaneous genetic/epigenetic haplotyping, as well as ultrasensitive detection of epigenetic transformations.

## ONLINE METHODS

### Human Subjects

This study was approved by The Johns Hopkins School of Medicine Institutional Review Board. The donor with a clinical diagnosis of FSHD 1 was confirmed by the University of Iowa Diagnostic Laboratories to have a contracted D4Z4 array on a 4qA allele by pulse-field gel electrophoresis and Southern blotting. The EcoRI/BlnI allele of 15 kb on chromosome 4qA corresponds to approximately 4-5 repeats, and of 27 kb allele on chromosome 4qB corresponds to approximately 8 repeats.

### DNA samples

λ-Bacteriophage DNA (New England BioLabs, Ipswich MA, USA) was used as provided. BAC DNA was purified from *E.coli* cells containing the CH16-291A23 BAC. Cells were cultured overnight in LB containing 12.5 μg/ml cloramphenicol (Sigma-Aldrich, Rehovot, Israel) at 30°C. BAC DNA was purified from the cells using the NucleoBondXtra BAC kit (MACHEREY-NAGEL Inc. Dü ren, Germany). DNA from a lymphocyte cell line (NA12878, Coriell Institute for Medical Research, Camden New Jersey, USA) and DNA from primary blood cells (purchased from HemaCare Inc. Van Nuys, CA, USA) were purified in agarose plugs in order to maintain large DNA fragments, following the Bionano Genomics IrysPrep protocol^29^.

### M.TaqI - assisted labeling for R^2^OM^2^

To generate methylation sensitive labels we used the DNA MTase M.TaqI, which catalyzes the transfer of a carboxytetramethylrhodamine (TAMRA) fluorophore from the synthetic cofactor AdoYnTAMRA onto the adenine residue within its recognition sequence (TCGA)^39^. The labeling reaction was carried out as follows: 1 μg of DNA was reacted with 37.5 ng of M.TaqI and 40 μM of AdoYnTAMRA in labeling buffer (20 mM Tris-HOAc, 10 mM Mg(Cl)2, 50 mM KOAc, 1 mM DTT, pH 7.9) in the presence of 0.01% Triton-X 100 and 0.1 mg/ml BSA in a total reaction volume of 25 μL at 60°C for 1 hour. The labeled DNA was reacted with 40 μg of proteinase K (Sigma-Aldrich, Rehovot, Israel) at 45°C for 1 hour to disassemble protein-DNA aggregates. For methylated samples CpG methylation was performed prior to labeling using the CpG-specific DNA MTase M.SssI (Thermo Scientific, Waltham, MA, USA) according to manufacturer’s instructions but with twice the suggested amount of enzyme to ensure complete methylation. To obtain partial methylation, the reaction was carried out using the recommended amount of enzyme, but for 75% of the recommended incubation time. Methylation was verified by digestion with a methylation-sensitive restriction enzyme HpaII (New England BioLabs Inc., Ipswich MA, USA), followed by gel electrophoresis to ensure that the DNA was protected or partially protected from restriction.

### Nick labeling for optical genome mapping

DNA was prepared in a nick-labeling-repair reaction (NLR) involves (a) the nicking enzyme Nb.BsmI, Nb.BssSI or Nt.BspQI which generates single-strand nicks at its specific recognition sites (GAATGC, CACGAG or GCTCTTCN respectively), (b) a DNA polymerase enzyme incorporates fluorescent nucleotides at the nicked sites; and finally, (c) a DNA ligase enzyme repairs the remaining single-strand breaks. For the NLR reaction involving Nb.BssSI or Nt.BspQI, DNA was labeled using the IrysPrep kit (Bionano Genomics Inc., San Diego CA, USA) according to manufacturer’s instructions. For Nb.BsmI-based NLR, DNA (900 ng) was first reacted with 30 units of the enzyme (New England BioLabs Inc., Ipswich MA, USA) in 30 μl NEBuffer 3.1 for 120 minutes at 65°C. Next, the DNA was reacted with 15 units of Taq DNA polymerase (New England BioLabs Inc., Ipswich MA, USA) in the presence of the following nucleotides: dGTP, dCTP dATP (Sigma-Aldrich, Rehovot, Israel) and the fluorescent nucleotide dUTP-Atto647 (Jena Bioscience GmbH, Jena, Germany) at a final concentration of 600 nM (each). The reaction was carried out in a reaction buffer (ThermoPol buffer, New England BioLabs Inc., Ipswich MA, USA) in a total volume of 45 μl for 60 min at 72°C. Finally, the DNA was reacted with 120 units of Taq DNA ligase (New England BioLabs Inc., Ipswich MA, USA) with 0.5 mM NAD+ (New England BioLabs Inc., Ipswich MA, USA), in a reaction buffer (ThermoPol Buffer, New England BioLabs Inc., Ipswich MA, USA) including 0.5 mM NAD+ (New England BioLabs Inc., Ipswich MA, USA) and of 10 μM dNTP mix, in a total reaction volume of 60 μL for 30 min at 45°C. For R^2^OM^2^ experiments, DNA was initially labeled with by NLR and then 0.05 μg - 0.5 μg of the labeled DNA was reacted with M.TaqI as described above (the reaction was scaled down accordingly).

### Sample preparation, DNA stretching and imaging

Post labeling, BAC and λ DNA were cleaned by ethanol precipitation as has been described previously^39^. Genomic DNA was cleaned by embedding it into agarose plugs and washing these in TE buffer (see supporting information). Prior to imaging, the labeled DNA was stained with 0.5 pμM of YOYO-1 (Invitrogen, Carlsbad, CA, USA) for visualization of its contour. DTT (Sigma-Aldrich, Rehovot, Israel) was added to the reaction (200 mM) to prevent photo bleaching and DNA breaks. To stretch the DNA from its random coil conformation into a linear form, allowing imaging of its contour, we used two types of experimental scheme: The first type of stretch used modified glass surfaces. In this approach, DNA sample-solutions were flowed by applying capillary forces or using microfluidics, on glass surfaces that were chemically modified to facilitate the DNA’s anchoring and stretching on the surfaces (see supporting information). After stretching, the DNA was imaged using an epifluorescence microscope (FEI Munich GmbH, Germany) equipped with a high-resolution EMCCD IXon888 camera (Andor Technology Ltd. Belfast, UK). A 150 W Xenon lamp (FEI Munich GmbH, Germany) was used for excitation with filter sets of 485/20ex and 525/30em, 560/25ex and 607/36em, and 650/13ex and 684/24em (Semrock Inc., Rochester NY, USA) for the YOYO-1, TAMRA, and Atto-647 channels respectively. For high-throughput mapping experiments, samples were analyzed in nanochannels array chips (Bionano Genomics Inc., San Diego CA, USA). On the chip, DNA is forced into 45 nm square nanochannels using an electric field and is stretched along the channel axis for imaging. This process is carried out in automated cycles by the Irys instrument (Bionano Genomics Inc., San Diego CA, USA) ^42^.

### Data analysis

Images of DNA molecules stretched on modified glass surfaces were manually aligned, according to the barcode created outside the repetitive region. This enabled detection of the starting point of the repeat array, allowing counting of individual repeat units.

For analysis of the high-throughput nanochannel array data, raw images were processed and labeled DNA molecules were detected and digitized by custom image-processing and analysis software (IrysView^29^). Genetic labels were assigned one set of coordinates along the molecules (the genetic map) and the methylation labels are assigned another set (the methylation map). The detection process output the number of labels per 100 kbp allowing for direct comparison between labeling level of different samples, and quantitative assessment of methylation levels. For R^2^OM^2^, DNA molecules are first mapped to the genome reference based on the match between the fluorescent genetic pattern along the molecule and the pattern expected from the known reference sequence. after alignment, the methylation maps reflected the distribution of non-methylated CpGs along the mapped genomic regions, and methylation profiles could be aligned to UCSC gene tracks ^53^ (https://genome.ucsc.edu/index.html). IrysView also enabled *de-novo* assembly of the detected molecules into a consensus barcode or contig (genome map) and its comparison with the theoretical barcode. This unsupervised assembly was used to assess the number of repeats in the FSHD BAC model system.

In order to generate plots of labeling intensity along the BAC sequence (Figure 4c), M.TaqI positions were extracted by extrapolation (in-house software) after molecule alignment facilitated by IrysView. We then counted the total number of M.TaqI labels at each genomic position using BEDTools^54^, as well as the total number of molecules aligned to each position. A score for each genomic position was calculated by dividing the total number of labels, attributed to it, by the total number of aligned molecules (simply, a score of 1 would mean that all the molecules overlapping this bp had a label at this bp, while 0 would mean that none of the molecules had such a label). This value was then normalized according to the minimum and maximum values of the data.

### Next-generation sequencing

Purified BAC DNA was sheared using Covaris AFA (Covaris Inc. Woburn MA, USA) and the fragments were size-separated by electrophoresis using agarose gel, allowing selective extraction of fragments within the range of 150-300 bp. Illumina sequencing libraries were prepared using NEXTFlex kit (Bioo Scientific Corporation, Austin TX, USA) and sequenced using MiSeq (Illumina Inc. San Diego, CA. USA) to a paired-end coverage of 15,000X. Sequencing reads were *de-novo* assembled using CLC Workbench software (CLC Bio-Qiagen, Aarhus, Denmark).

## ACKNOWLEDGEMENTS

We thank Prof. D. Gabellini for his kind gift of the CH16-291A23 BAC and K. Glensk for preparing M.TaqI. We acknowledge financial support from the German-Israeli foundation [I-1196-195.9/2012] (E.W. and Y.E.), the BeyondSeq consortium [EC program 63489] (Y.E.), the European Research Councils starter grant [337830] (Y.E.) i-Core program of the Israel Science Foundation [1902/12] ( Y.E.), NIH U54HD060848 (KRW), and NIH U01 MH106884 (JP). We thank Kerstin Glensk for preparing M.TaqI.

## REFERENCES

(1) Bird, A. DNA Methylation Patterns and Epigenetic Memory. Genes Dev. 2002, 16, 6–21.

(2) Jabbari, K.; Bernardi, G. Cytosine Methylation and CpG, TpG (CpA) and TpA Frequencies. Gene 2004, 333, 143–149.

(3) Esteller, M. CpG Island Hypermethylation and Tumor Suppressor Genes: A Booming Present, a Brighter Future. Oncogene 2002, 21, 5427–5440.

(4) Jones, P. A. Functions of DNA Methylation: Islands, Start Sites, Gene Bodies and Beyond. Nat. Rev. Genet. 2012, 13, 484–492.

(5) Schmid, C. W. Organization of the Human Genome Transcription. 345–358.

(6) Batzer, M. a; Deininger, P. L. Alu Repeats and Human Genomic Diversity. Nat. Rev. Genet. 2002, 3, 370–379.

(7) Mather, K. A.; Jorm, A. F.; Parslow, R. A.; Christensen, H. Is Telomere Length a Biomarker of Aging? A Review. J. Gerontol.A. Biol. Sci. Med. Sci. 2011, 66, 202–213.

(8) M. Duyao, C. Ambrose,R. Myers,A. Novelletto,F. Persichetti,M. Frontali,S. Folstein,C. Ross,M. Franz,M. Abbott,J. Gray,P. Conneally,A. Young,J. Penney,Z. Hollingsworth,I. Shoulson,A. Lazzarini,A. Falek,W. Koroshetz,D. Sax,E. Bird,J. Von, & M.M. Trinucleotide Repeatlength Instability and Age of Onset in Huntington’s Disease. Nat. Genet. 1993, 4, 387–392.

(9) Edwards, A.; Hammond, H. A.; Jin, L.; Caskey, C. T.; Chakraborty, R. Genetic Variation at Five Trimeric and Tetrameric Tandem Repeat Loci in Four Human Population Groups. Genomics 1992, 12, 241–253.

(10) Maarel, M. Van Der; Deidda, G.; Lemmers, R. J. L. F.; Overveld, P. G. M. Van; Wielen, M. Van Der; Hewitt, J. E.; Sandkuijl, L.; Bakker, B.; Ommen, G. B. Van; Padberg, G. W.; et al. De Novo Facioscapulohumeral Muscular Dystrophy: Frequent Somatic Mosaicism, Sex-Dependent Phenotype, and the Role of Mitotic Transchromosomal Repeat Interaction between Chromosomes 4 and 10. 1996, 4, 26–35.

(11) Chamberlain, N. L.; Driver, E. D.; Miesfeld, R. L. The Length and Location of CAG Trinucleotide Repeats in the Androgen Receptor N-Terminal Domain Affect Transactivation Function. Nucleic Acids Res. 1994, 22, 3181–3186.

(12) Balog, J.; Miller, D.; Sanchez-Curtailles, E.; Carbo-Marques, J.; Block, G.; Potman, M.; de Knijff, P.; Lemmers, R. J. L. F.; Tapscott, S. J.; van der Maarel, S. M. Epigenetic Regulation of the X-Chromosomal Macrosatellite Repeat Encoding for the Cancer/testis Gene CT47. Eur.J. Hum. Genet. 2012, 20, 185–191.

(13) Pook, M. A. DNA Methylation and Trinucleotide Repeat Expansion Diseases. In DNA Methylation - From Genomics to Technology. InTech, 2012.

(14) Hansen, K. D.; Timp, W.; Bravo, H. C.; Sabunciyan, S.; Langmead, B.; McDonald, O. G.; Wen, B.; Wu, H.; Liu, Y.; Diep, D.; et al. Increased Methylation Variation in Epigenetic Domains across Cancer Types. Nat. Genet. 2011, 43, 768–775.

(15) Huichalaf, C.; Micheloni, S.; Ferri, G.; Caccia, R.; Gabellini, D. DNA Methylation Analysis of the Macrosatellite Repeat Associated with FSHD Muscular Dystrophy at Single Nucleotide Level. PLoS One 2014, 9, e115278.

(16) Gaillard, M.-C.; Roche, S.; Dion, C.; Tasmadjian, A.; Bouget, G.; Salort-Campana, E.; Vovan, C.; Chaix, C.; Broucqsault, N.; Morere, J.; et al. Differential DNA Methylation of the D4Z4 Repeat in Patients with FSHD and Asymptomatic Carriers. Neurology 2014, 83, 733–742.

(17) Sacconi, S.; Lemmers, R. J. L. F.; Balog, J.; van der Vliet, P. J.; Lahaut, P.; van Nieuwenhuizen, M. P.; Straasheijm, K. R.; Debipersad, R. D.; Vos-Versteeg, M.; Salviati, L.; et al. The FSHD2 Gene SMCHD1 Is a Modifier of Disease Severity in Families Affected by FSHD1. Am.J. Hum. Genet. 2013, 93, 744–751.

(18) INSERM and French Ministry of Health. Prevalence of Rare Diseases: Bibliographic Data http://www.orpha.net/orphacom/cahiers/docs/GB/Prevalence_of_rare_diseases_by_alphabetical_list.pdf.

(19) Korshunova, Y.; Maloney, R. K.; Lakey, N.; Citek, R. W.; Bacher, B.; Budiman, A.; Ordway, J. M.; McCombie, W. R.; Leon, J.; Jeddeloh, J. A.; et al. Massively Parallel Bisulphite Pyrosequencing Reveals the Molecular Complexity of Breast Cancer-Associated Cytosine-Methylation Patterns Obtained from Tissue and Serum DNA. Genome Res. 2008, 18, 19–29.

(20) Gu, H.; Smith, Z. D.; Bock, C.; Boyle, P.; Gnirke, A.; Meissner, A. Preparation of Reduced Representation Bisulfite Sequencing Libraries for Genome-Scale DNA Methylation Profiling. Nat. Protoc. 2011, 6, 468–481.

(21) Guo, H.; Zhu, P.; Wu, X.; Li, X.; Wen, L.; Tang, F. Single-Cell Methylome Landscapes of Mouse Embryonic Stem Cells and Early Embryos Analyzed Using Reduced Representation Bisulfite Sequencing. 2013, 2126–2135.

(22) Nagano, T.; Lubling, Y.; Stevens, T. J.; Schoenfelder, S.; Yaffe, E.; Dean, W.; Laue, E. D.; Tanay, A.; Fraser, P. Single-Cell Hi-C Reveals Cell-to-Cell Variability in Chromosome Structure. Nature 2013, 502, 59–64.

(23) Treangen, T. J.; Salzberg, S. L. Repetitive DNA and next-Generation Sequencing: Computational Challenges and Solutions. Nat. Rev. Genet. 2012, 13, 36–46.

(24) Vranken, C.; Deen, J.; Dirix, L.; Stakenborg, T.; Dehaen, W.; Leen, V.; Hofkens, J.; Neely, R. K. Super-Resolution Optical DNA Mapping via DNA Methyltransferase-Directed Click Chemistry. Nucleic Acids Res. 2014, 42, e50.

(25) Nilsson, A. N.; Emilsson, G.; Nyberg, L. K.; Noble, C.; Svensson Stadler, L.; Fritzsche, J.; Moore, E. R. B.; Tegenfeldt, J. O.; Ambjörnsson, T.; Westerlund, F. Competitive Binding-Based Optical DNA Mapping for Fast Identification of Bacteria - Multi-Ligand Transfer Matrix Theory and Experimental Applications on Escherichia Coli. Nucleic Acids Res. 2014, 42, E118.

(26) Meng, X.; Benson, K.; Chada, K.; Huff, E. J.; Schwartz, D. C. Optical Mapping of Lambda Bacteriophage Clones Using Restriction Endonucleases. Nat. Genet. 1995, 9, 432–438.

(27) Levy-Sakin, M.; Ebenstein, Y. Beyond Sequencing: Optical Mapping of DNA in the Age of Nanotechnology and Nanoscopy. Curr. Opin. Biotechnol. 2013, 1–9.

(28) Levy-Sakin, M.; Grunwald, A.; Kim, S.; Gassman, N. R.; Gottfried, A.; Antelman, J.; Kim, Y.; Ho, S. O.; Samuel, R.; Michalet, X.; et al. Toward Single-Molecule Optical Mapping of the Epigenome. ACS Nano 2014, 8, 14–26.

(29) Cao, H.; Hastie, A. R.; Cao, D.; Lam, E. T.; Sun, Y.; Huang, H.; Liu, X.; Lin, L.; Andrews, W.; Chan, S.; et al. Rapid Detection of Structural Variation in a Human Genome Using Nanochannel-Based Genome Mapping Technology. Gigascience 2014, 3, 1–11.

(30) Mostovoy, Y.; Levy-Sakin, M.; Lam, J.; Lam, E. T.; Hastie, A. R.; Marks, P.; Lee, J.; Chu, C.; Lin, C.; Džakula, Ž.; et al. A Hybrid Approach for de Novo Human Genome Sequence Assembly and Phasing. Nat. Methods 2016, 12–17.

(31) Nifker, G.; Levy-Sakin, M.; Berkov-Zrihen, Y.; Shahal, T.; Gabrieli, T.; Fridman, M.; Ebenstein, Y. One-Pot Chemoenzymatic Cascade for Labeling of the Epigenetic Marker 5-Hydroxymethylcytosine. Chembiochem 2015, 16, 1857–1860.

(32) Zirkin, S.; Fishman, S.; Sharim, H.; Michaeli, Y.; Don, J.; Ebenstein, Y. Lighting up Individual DNA Damage Sites by in Vitro Repair Synthesis. J. Am. Chem. Soc. 2014, 136, 7771–7776.

(33) Michaeli, Y.; Shahal, T.; Torchinsky, D.; Grunwald, A.; Hoch, R.; Ebenstein, Y. Optical Detection of Epigenetic Marks: Sensitive Quantification and Direct Imaging of Individual Hydroxymethylcytosine Bases. Chem. Commun. (Camb). 2013, 49, 8599–8601.

(34) Gilboa, T.; Torfstein, C.; Juhasz, M.; Grunwald, A.; Ebenstein, Y.; Weinhold, E.; Meller, A. Single-Molecule DNA Methylation Quantification Using Electro-Optical Sensing in Solid-State Nanopores. ACS Nano 2016, 10, 8861–8870.

(35) GeneCards - Human Genes | Gene Database | Gene Search http://www.genecards.org/ (accessed Jan 8, 2017).

(36) Kinkley, S.; Staege, H.; Mohrmann, G.; Rohaly, G.; Schaub, T.; Kremmer, E.; Winterpacht, A.; Will, H. SPOC1: A Novel PHD-Containing Protein Modulating Chromatin Structure and Mitotic Chromosome Condensation. J. Cell Sci. 2009, 122.

(37) Hanz, G. M.; Jung, B.; Giesbertz, A.; Juhasz, M.; Weinhold, E. Sequence-Specific Labeling of Nucleic Acids and Proteins with Methyltransferases and Cofactor Analogues. J. Vis. Exp. 2014, 3–12.

(38) Klimasauskas, S.; Weinhold, E. A New Tool for Biotechnology: AdoMet-Dependent Methyltransferases. Trends Biotechnol. 2007, 25, 99–104.

(39) Grunwald, A.; Dahan, M.; Giesbertz, A.; Nilsson, A.; Nyberg, L. K.; Weinhold, E.; Ambjörnsson,, T.; Westerlund, F.; Ebenstein, Y. Bacteriophage Strain Typing by Rapid Single Molecule Analysis. Nucleic Acids Res. 2015, 1–8.

(40) McClelland, M.; Nelson,M. Effect of Site-Specific Méthylation on Dna Modification Methyltransferases and Restriction Endonucleases. Nucleic Acids Res. 1992, 20, 2145–2157.

(41) Hansen, K. D.; Sabunciyan, S.; Langmead, B.; Nagy, N.; Curley, R.; Klein, G.; Klein, E.; Salamon, D.; Feinberg, A. P. Large-Scale Hypomethylated Blocks Associated with Epstein-Barr Virus – Induced B-Cell Immortalization. 2014, 177–184.

(42) Das, S. K.; Austin, M. D.; Akana, M. C.; Deshpande, P.; Cao, H.; Xiao, M. Single Molecule Linear Analysis of DNA in Nano-Channel Labeled with Sequence Specific Fluorescent Probes. Nucleic Acids Res. 2010, 38, e177.

(43) Bernstein, B. E.; Stamatoyannopoulos, J. A.; Costello, J. F.; Ren, B.; Milosavljevic, A.; Meissner, A.; Kellis, M.; Marra, M. A.; Arthur, L.; Ecker, J. R.; et al. The NIH Roadmap Epigenomics Mapping Consortium. NIH Public Access 2013, 28, 1045–1048.

(44) Reinius, L. E.; Acevedo, N.; Joerink, M.; Pershagen, G.; Dahlén, S. E.; Greco, D.; Söderhäll, C.; Scheynius, A.; Kere, J. Differential DNA Methylation in Purified Human Blood Cells: Implications for Cell Lineage and Studies on Disease Susceptibility. PLoS One 2012, 7.

(45) Cabianca, D. S.; Gabellini, D. The Cell Biology of Disease: FSHD: Copy Number Variations on the Theme of Muscular Dystrophy. J. Cell Biol. 2010, 191, 1049–1060.

(46) Yoon, S.; Xuan, Z.; Makarov, V.; Ye, K.; Sebat, J. Sensitive and Accurate Detection of Copy Number Variants Using Read Depth of Coverage. Genome Res. 2009, 19, 1586–1592.

(47) Thomas Anantharaman, B. M. False Positives in Genomic Map Assembly and Sequence Validation file:///C:/Users/grunwald/Downloads/FalsePositives.pdf (accessed Mar 30, 2016).

(48) Pendleton, M.; Sebra, R.; Pang, A. W.C.; Ummat, A.; Franzen, O.; Rausch, T.; Stütz, A. M.; Stedman, W.; Anantharaman, T.; Hastie, A.; et al. Assembly and Diploid Architecture of an Individual Human Genome via Single-Molecule Technologies. Nat. Methods 2015, 12, 780–786.

(49) Jeffet, J.; Kobo, A.; Su, T.; Grunwald, A.; Green, O.; Nilsson, A. N.; Eisenberg, E.; Ambjörnsson, T.; Westerlund, F.; Weinhold, E.; et al. Super-Resolution Genome Mapping in Silicon Nanochannels. ACS Nano 2016, 10, 9823–9830.

(50) Lemmers, R. J. L. F.; Wohlgemuth, M.; van der Gaag, K. J.; van der Vliet, P. J.; van Teijlingen, C. M. M.; de Knijff, P.; Padberg, G. W.; Frants, R. R.; van der Maarel, S. M. Specific Sequence Variations within the 4q35 Region Are Associated with Facioscapulohumeral Muscular Dystrophy. Am. J. Hum. Genet. 2007, 81, 884–894.

(51) Lemmers, R. J. L. F.; Tawil, R.; Petek, L. M.; Balog, J.; Block, G. J.; Santen, G. W. E.; Amell, A. M.; van der Vliet, P. J.; Almomani, R.; Straasheijm, K. R.; et al. Digenic Inheritance of an SMCHD1 Mutation and an FSHD-Permissive D4Z4 Allele Causes Facioscapulohumeral Muscular Dystrophy Type 2. Nat. Genet. 2012, 44, 1370–1374.

(52) Choi, S. H.; Worswick, S.; Byun, H.-M.; Shear, T.; Soussa, J. C.; Wolff, E. M.; Douer, D.; Garcia-Manero, G.; Liang, G.; Yang, A. S. Changes in DNA Methylation of Tandem DNA Repeats Are Different from Interspersed Repeats in Cancer. Int. J. Cancer 2009, 125, 723–729.

(53) Kent, W. J.; Sugnet, C. W.; Furey, T. S.; Roskin, K. M. The Human Genome Browser at UCSC W. J. Med. Chem. 1976, 19, 1228–1231.

(54) Quinlan, A. R.; Hall, I. M. BEDTools: A Flexible Suite of Utilities for Comparing Genomic Features. Bioinformatics 2010, 26, 841–842.

